# Depth and location influence prokaryotic and eukaryotic microbial community structure in New Zealand fjords

**DOI:** 10.1101/680694

**Authors:** Sven P. Tobias-Hünefeldt, Stephen R. Wing, Nadjejda Espinel-Velasco, Federico Baltar, Sergio E. Morales

## Abstract

Systems with strong horizontal and vertical gradients, such as fjords, are useful models for studying environmental forcing. Here we examine microbial (prokaryotic and eukaryotic) community changes associated with the surface low salinity layer (LSL) and underlying seawater in multiple fjords in Fiordland National Park (New Zealand). High rainfall (1200-8000 mm annually) and linked runoff from native forested catchments results in surface LSLs with high tannin concentrations within each fjord. These gradients are expected to drive changes in microbial communities. We used amplicon sequencing (16S and 18S) to assess the impact of these gradients on microbial communities and identified depth linked changes in diversity and community structure. With increasing depth we observed significant increases in Proteobacteria (15%) and SAR (37%), decreases in Opisthokonta (35%), and transiently increased Bacteroidetes (3% increase from 0 to 40 m, decreasing by 8% at 200 m). Community structure differences were observed along a transect from inner to outer regions, specifically 25% mean relative abundance decreases in Opisthokonta and Bacteroidetes, and increases in SAR (25%) and Proteobacteria (>5%) at the surface, indicating changes based on distance from the ocean. This provides the first in-depth view into the ecological drivers of microbial communities within New Zealand fjords.

## 1. Introduction

Microorganisms play a major role in the cycling of both organic and inorganic nutrients in all major ecosystems (Falkowski et al., 2008; Junge et al., 2006; Sullam et al., 2017; Zahran, 1999). In particular, marine microbes have been linked to global nutrient cycling with community changes affecting global balances of carbon and nitrogen (Arrigo, 2005). Further, as both primary producers and recyclers, they provide information about ecosystem health, viability, and function (Azam and Worden, 2004; Graham et al., 2016). It is therefore critical to understand the factors controlling the microbial community structure in the marine environments. Up to date, a wide range of marine ecosystems remains understudied, including nearshore systems such as fjords. Indeed, fjord-systems with strong horizontal and vertical environmental gradients can be extremely useful models for studying the response of microbial communities to environmental factors.

Fjords are long narrow coastal inlets flanked by steep cliffs, typically carved by glaciers and resulting in deep waterways. In New Zealand (NZ), these sites receive large amounts of rain (>400 mm per month) (NIWA, 2016) and carbon per unit area (Smith et al., 2015) through exogenous carbon inputs particularly tannins (Currey et al., 2009). The external freshwater inputs help create vertical and horizontal gradients along the transition towards inland from the sea, and with depth, consistent with other sites (Storesund et al., 2017). These gradients can differ based on geographic (i.e. catchment area size) and environmental (i.e. land cover) variables resulting in highly diverse marine, and microbial communities, even in relatively close geographical locations (Brattegard, 1980; Herlemann et al., 2011). Microbial marine communities are dynamic (Berdjeb et al., 2018; Henriques et al., 2006) arising from the temporally and spatially fluid aquatic gradients (Jakacki et al., 2017; Storesund et al., 2017). Despite this, the microbial communities within fjords of the southern hemisphere (particularly NZ) remain unexplored. NZ fjords are part of a temperate climate, unlike those found in Chile, Antarctica and Iceland. The different climates, fauna and flora changes, as well as morphological differences, such as depth, could influence the drivers of microbial community composition in fjords.

In the present study we characterised the microbial (both eukaryotic and prokaryotic) community composition of six fjords in New Zealand, and linked the observed variance in diversity and community composition to key environmental variables (i.e. depth, salinity, horizontal location, oxygen, temperature) (Logue et al., 2015). We hypothesised that, similar to other fjord systems around the world, depth and salinity would be key determinants of microbial diversity and community composition (Sakami et al., 2016; Storesund et al., 2017; Walsh et al., 2016; Ying et al., 2009). We also hypothesised a close coupling between changes in eukaryotic and prokaryotic communities shift along fjords. To test this, we collected samples from six New Zealand fjords at multiple depths and locations, utilising amplicon sequencing of the small subunit ribosomal rRNA gene (both 16S and 18S) to determine changes in microbial communities.

## 2. Materials and methods

### 2.1. Sampling

Samples were collected in November 2015 throughout Fiordland National Park (45.60° S, 167.36° E) (Fig. 1), specifically Breaksea Sound (45.5860° S, 166.7567° E), Chalky Inlet (45.9922° S, 166.6024° E), Doubtful Sound (45.3260° S, 166.9911° E), Dusky Sound (45.7246° S, 166.5065° E), Long Sound (46.0001° S, 166.762415° E), and Wet Jacket Arm (45.6415° S, 166.8413° E).

**Figure 1.**
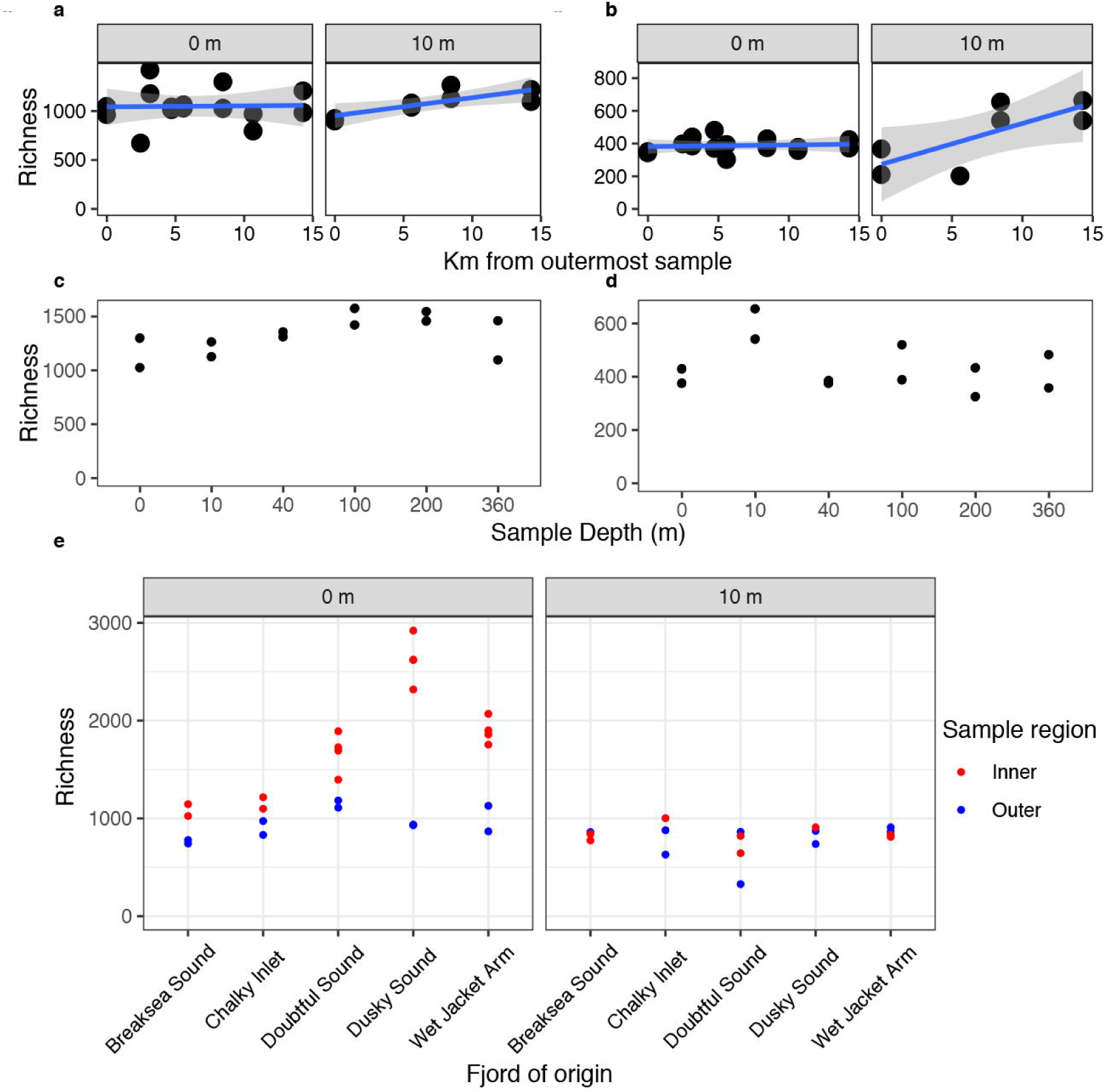
Geographical location of sampling sites. Geographical locations of sampling sites in New Zealand, Fiordland (a) (Kahle and Wickham, 2013), and Long Sound’s sampling scheme (b).

A total of 44 samples were collected from Breaksea Sound (8), Chalky Inlet (10), Doubtful Sound (10), Dusky Sound (10), and Wet Jacket Arm (6). At each site duplicate (2) samples were collected from inner and outer regions of the fjord, and for each region sampling was performed at two depths (0 and 10 m), exceptions including Breaksea Sound’s outer 10 m, Chalky Inlet’s inner 10 m, Doubtful Sound’s inner surface, Dusky Sound’s inner 10 m, and Wet Jacket Arm’s inner surface regions where 1, 1, 4, 1, and 4 samples were taken respectively. For higher resolution sampling, duplicate samples were collected at Long Sound from a transect starting at an outer location (Fig. 1 b) and moving inwards with sampling occurring at 2.47, 3.16, 4.73, 5.59, 8.47, 10.67, and 14.3 km from the outermost sample at a depth of 0 and 10 m, exceptions including samples at 2.47, 3.16, 4.73, and 10.67 km at 10 m where no samples were taken, at 8.47 km where only a single sample was taken along the surface for prokaryotes, and at 10.67 km at the surface where only a single eukaryotic sample was taken. An additional depth profile was collected at 8.47 km from the outermost site with duplicate samples collected at 0, 10, 40, 100, 200, and 360 m depths (Fig. 1 b), exceptions including surface, 10 m, and 200 m samples were 4, 3, and 3 samples were taken respectively.

A CTD sensor system (SBE-25 0352) was used for vertical profiling of salinity (conductivity), temperature, and dissolved oxygen. Water samples were collected in 10 L Niskin bottles for subsequent analysis.

### 2.2. DNA extraction

A 0.5 to 1 litre subsample of water was filtered through a 0.22 µm (diameter of 47 mm) polycarbonate filter prior to freezing and storage at −20 °C during transport to the lab and finally stored at −80°C until further processing. Total community DNA was extracted from filters using the MoBio DNeasy® PowerSoil® Kit (MoBio, Solana Beach, CA, USA) in accordance to a modified manufacturer’s protocol. Samples were bead beaten in a Geno/Grinder for 2 × 15s instead of vortexing at maximum speed for 10 minutes. All extracted DNA was stored at −20°C until further use.

### 2.3. Total community profiling using small subunit ribosomal rRNA gene (SSU rRNA) amplicon libraries

Community profiles were generated using barcoded 16S (targeting the V4 region: 515F (5’-NNNNNNNNGTGTGCCAGCMGCCGCGGTAA-3’) and 806R (5’-GGACTACHVGGGTWTCTAAT-3’)) or 18S 1391f (5’-GTACACACCGCCCGTC-3’) and EukBr (5’-TGATCCTTCTGCAGGTTCACCTAC-3’) rRNA gene primers as per the Earth Microbiome Project protocol(Caporaso et al., 2012). Barcoded samples were then loaded onto separate Illumina HiSeq (16S) or MiSeq (18S) 2 × 151 bp runs (Illumina, Inc., CA, USA).

Raw community profiles were analysed using the Quantitative Insights Into Molecular Ecology (QIIME) 1.9.1 open-reference operation taxonomic unit-picking workflow (Caporaso et al., 2010). Default parameters, including a read length >75 bp, minimum number of consecutive high quality base calls to include a read as a fraction of the input read length of 0.75, a Phred quality score of 3, no ambiguous bases, and no mismatches in the primer sequence (Bokulich et al., 2013) were used. Analysed sequences were all >136 bp in length. Raw sequences were demultiplexed with no ambiguous bases, using only forward reads and the split_libraries_fastq.py command. This has been shown to increase sequence depth per sample and analysis speed while producing comparable results to paired end data (Werner et al., 2012). Operational taxonomic units (OTUs) were generated by clustering at 97% for 16S and 99% for 18S similarity using UCLUST (Edgar, 2010) and an open reference strategy based on reference sequences located in the SILVA database (release 128) using the QIIME pick_open_reference_otus.py command (Quast et al., 2013). OTUs were classified to 7 taxa levels (kingdom, phylum, class, order, family, genus, OTU) using BLAST (Altschul et al., 1997) with a maximum e-value of 0.001 against the SILVA database.

The 16S sequence pool was subsampled and rarefied 10 times to a depth of 22,000 sequences to eliminate biases in sampling depth, while the 18S sequence pool utilised a depth of 6,600 sequences. The 10 resulting tables were merged into a single OTU table with a total sequence depth of 220,000 and 66,000 sequences per sample respectively. The data was exported as a biom (json) file for further processing.

### 2.4. Microbial community analysis

All data analysis was carried out within RStudio (R Core Team, 2018) version 1.1.453 (RStudio Team, 2016), and visualised using the ggplot2 package (version 3.0.0) (Wickham, 2016) unless otherwise stated. Sample counts were transformed using transform_sample_counts() in the phyloseq package (version 1.24.2) (McMurdie and Holmes, 2013) to account for multiple rarefactions. Individual OTU abundances were divided by 10 to provide a mean. To avoid counts represented by fractions all data was rounded to the nearest possibility (using the round() command from the stats package – version 3.5.1) prior to downstream analysis. The α-diversity measures observed richness and Shannon were calculated using the estimate_richness() command within the phyloseq package, while Pielou evenness was calculated using the evenness() commands from the microbiome package (version 1.2.1) (Lahti et al., 2017). Significance was calculated using a Kruskal-Wallis (KW) test using the kruskal.test() command, from the stats package. Interactions between geographical parameters were calculated using the interaction() command from the base package. Visualisation of β-diversity utilised plot_ordination() from the vegan package together with adjustments from the ggplot2 package (version 3.0.0). Significant correlations between environmental variables and β-diversity were calculated using the PCA() command from the FactoMineR package (version 1.4.1) (Lê et al., 2008). A mantel test was used, as previously described, to identify significant community changes in relation to depth, horizontal location, and the fjord of origin.

Significant microbial community genera changes were calculated using the EdgeR package (version 3.22.3) (Robinson et al., 2009) and an exact test across both the five-fjord and Long Sound fjords. All p values were FDR adjusted using Bonferroni via the p.adjust() command from the stats package. All significantly changing organisms were displayed at the phyla level for ease of visualisation.

Medians were calculated using the ddply() command from the plyr package (version 1.8.4) (Wickham, 2011). Standard error, standard deviation and the 95% confidence interval calculated using summarySE() from Rmisc (version 1.5) (Hope, 2013), and arrange() from the dplyr package (version 0.7.6) (Wickham, 2018).

### 2.5. Statistical analyses

Physicochemical parameters across and within Long Sound were visualised using the graphics package (version 3.5.1) within RStudio (R Core Team, 2018). Mantel tests (vegan package version 2.5-2) (Oksanen, 2008), were used to determine significantly changing physicochemical parameters across geographical (depth, horizontal location, and fjord of origin) parameters.

### 2.6. Data availability

The sequence data from this study have been deposited in NCBI under BioProject PRJNA540153. All data generated and/or analysed during the study is available within the GitHub repository, https://github.com/SvenTobias-Hunefeldt/Fiordland_2019/.

## 3. Results

### 3.1. Alpha diversity within fjords

OTU richness remained constant across the surface transect in Long Sound fjord for both prokaryotic (1050 ± 183) and eukaryotic (388 ± 44) communities (Spearman, p value >0.05)(Fig. 2a-b). At 10 m depth, observed OTUs increased (Spearman: prokaryotes rho = 0.83, p value = 0.01; eukaryotes rho = 0.76, p value = 0.05) towards the head (landside/in) of the fjord resulting in a near doubling of richness (Fig. 2a-b) compared to the mouth of the fjord (seaside/out). On average, across the length of Long Sound, no significant differences (KW, p > 0.05) were detected in alpha diversity between surface and 10 m richness for either prokaryotes or eukaryotes.

**Figure 2.**
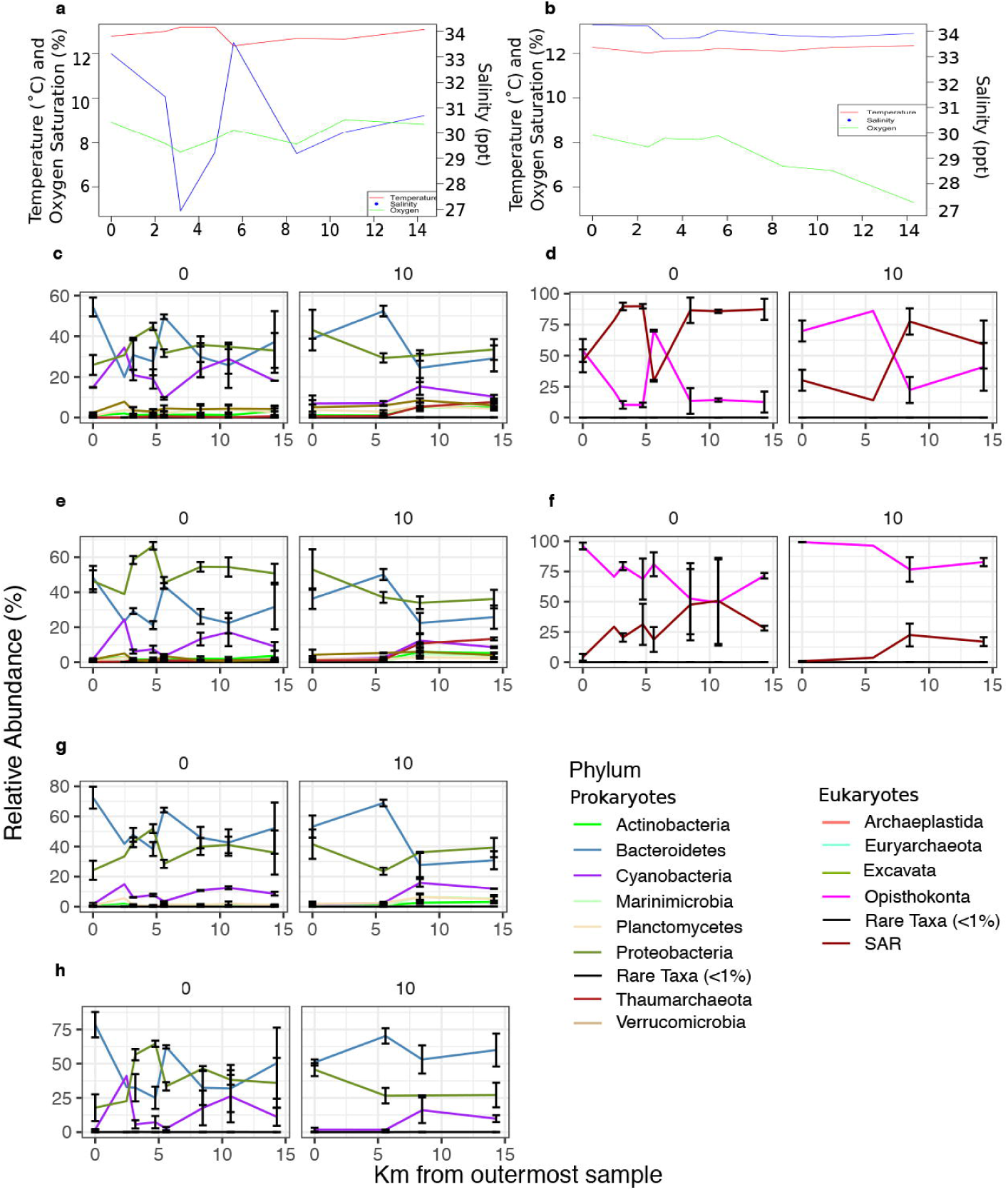
Alpha-diversity of Fiordland fjords. Richness for both 16S (left side) and 18S (right side) across Long Sound’s horizontal axis (a, b), Long Sound’s vertical axis (c, d), and richness based on 16S for all five sampled fjords samples (e). OTU clustering as carried out at 97% similarity.

A depth profile in the deepest sampled site of the same fjord (8.5 km) showed the prokaryotic richness increasing with depth, with a maximum observed (1503) at 100-200 m, whereas eukaryote’s highest richness (599) was located at 10 m (Fig. 2c-d). Overall half as many eukaryotes were identified in the sample compared to prokaryotes (Fig. 2).

When richness patterns between outer and inner sites were compared across 5 fjords (Breaksea Sound, Chalky Inlet, Doubtful Sound, Dusky Sound, and Wet Jacket Arm), inner surface communities exhibited higher prokaryotic richness (Fig. 2e), although high variance was observed across fjords. Both depth (KW, X^2^ of 19.55, p value < 0.01) and location within the fjord (KW, X^2^ of 9.40, p value < 0.01) were significantly associated with changes in observed richness, with strong interactions between them (KW, X^2^ of 29.75, p value < 0.01) (Table S1 and Fig. 2e).

### 3.2. Beta diversity within Long Sound

Long Sound’s microbial community was dominated by a few key phyla, with Proteobacteria and Bacteroidetes being the most abundant prokaryotes, while SAR (Stramenopila, Alveolate, and Rhizaria) and Opisthokonta were the most abundant eukaryotes (Table S2-S4). Changes in communities (β-diversity) within the same fjord where strongly linked to depth (Fig. 3). At Long Sound, samples along the horizontal transect were primarily clustered based on depth (0 vs. 10 m), for both prokaryotes (adonis (ado), r2 = 0.24, p value < 0.01, ANOSIM (ANO), R = 0.57, p value < 0.01) and eukaryotes (ado, r2 = 0.26, p value <0.01, ANO, R = 0.66, p value <0.01) (Fig. 3a-b), with interactions by location (Table S5). The same depth stratification was seen on the vertical axis of Long Sound, showing a clear difference between depths for eukaryotes (ado, r2 = 0.23, p value < 0.01, ANO, R = 1, p value <0.01), specifically between the surface, 10 m, and ≥40 m samples (Fig. 3c-d), but not prokaryotes (p value > 0.05). However, prokaryotic community changes were significantly associated with changes in salinity, oxygen and their interaction (Table S5), which were themselves associated with depth stratification (p value <0.05). For eukaryotic communities, stratification along the horizontal and vertical axis was correlated significantly with all tested environmental parameters excluding horizontal location (Table 14). Prokaryotic stratification based on depth was most correlated along the vertical NMDS axis (Fig. 3c-d), while the eukaryotic NMDS1 showed separation between communities above 10 and those below 40 m. The NMDS2 separated surface, 10 m, and below 40 m, the surface and 10 m communities shown to be more dissimilar than those below 40 m. Outer region prokaryotic and eukaryotic Long Sound samples clustered together, unlike the innermost samples (Fig. 3a-b). This pattern was also noted in the prokaryotic five-fjord NMDS (Fig. 4).

**Figure 3.**
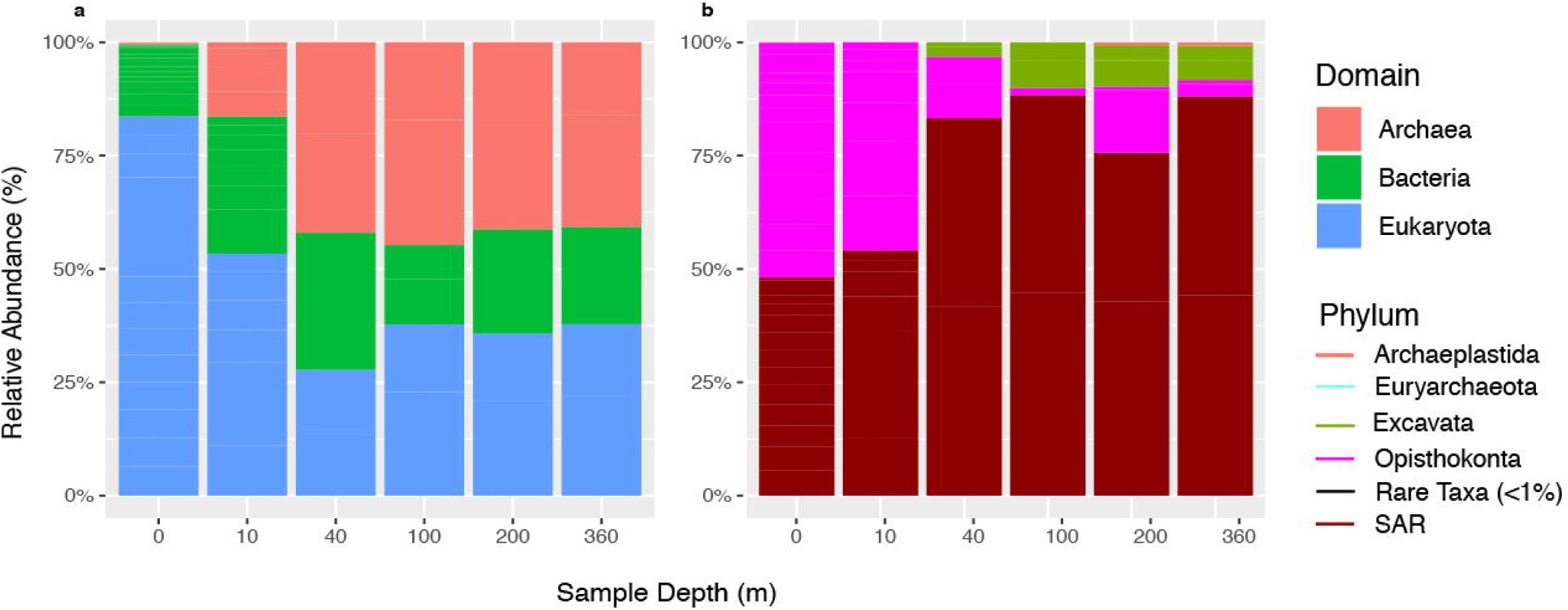
Microbial beta-diversity within Long Sound. Beta-diversity based on 16S (left panels) and 18S (right panels) data for Long Sound’s horizontal axis (a, b) and its vertical axis (c, d). Dissimilarity was assessed using Bray-Kurtis distance matrices based on OTUs at 97% similarity.

**Figure 4.**
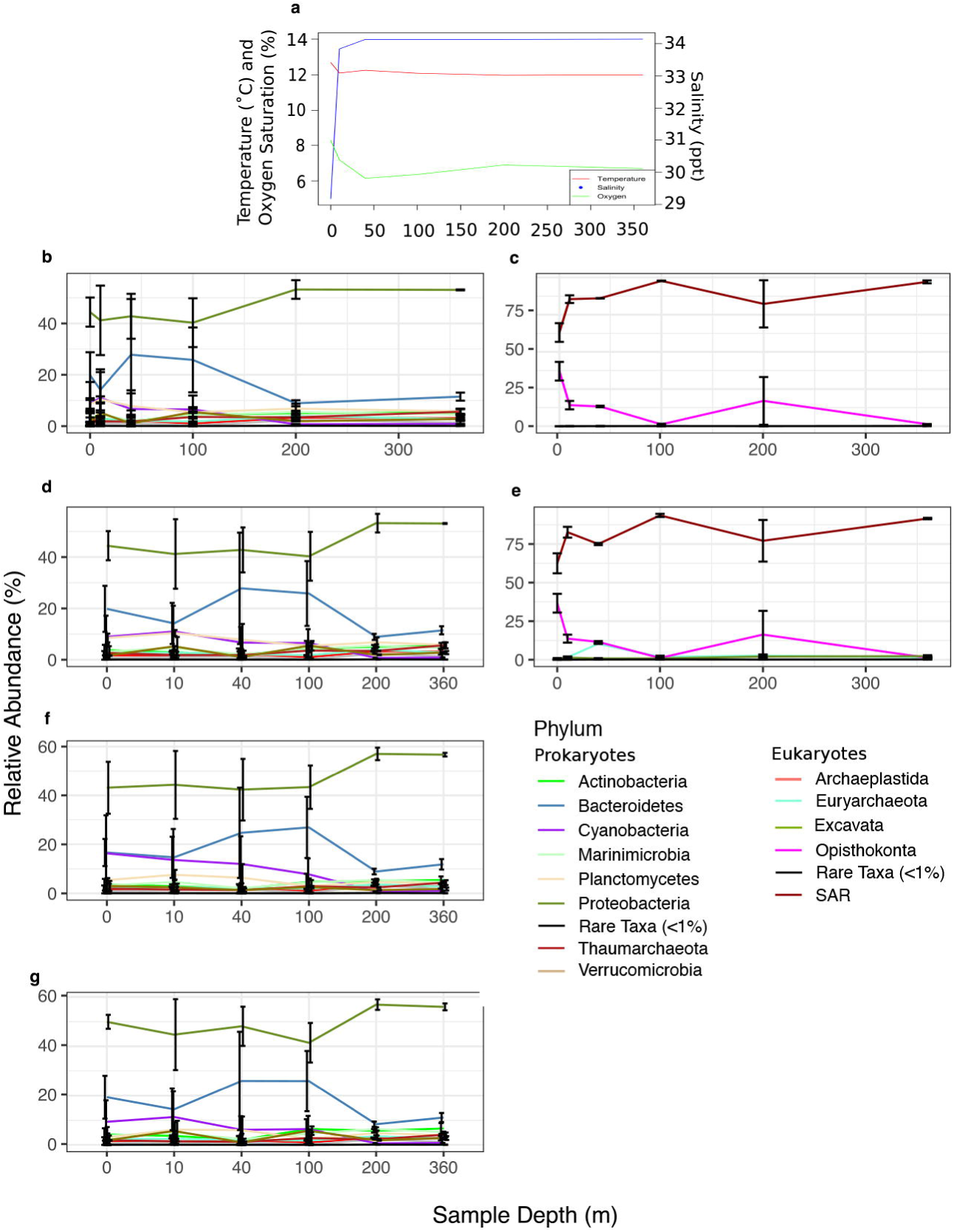
Microbial beta-diversity of five fjords. The fjord of origin, sample region, and depth were used to identify sample origin. Dissimilarity was assessed using Bray-Kurtis distance matrices based on OTUs at 97% similarity.

### 3.3. Significantly changing taxa

Salinity change patterns along the transect in deeper (10 m) samples are similar to the ones observed in surface samples, but less pronounced. Oxygen levels remained constant along the surface samples and showed a decrease towards the Fjord’s head in 10 m samples (Fig. 5a-b). The shifts found in community composition seemed to correlate with these changes. For example, along the transect, shifts in dominant taxa in the surface samples seemed to correlate to sharp changes in salinity. The most notable shift in taxa occurred between 5.6 and 8.5 km for both prokaryotes and eukaryotes, where the Bacteroidetes and Opisthokonta decreased from the outermost to innermost sample along a region with steep salinity changes (Fig. 5).

**Figure 5.**
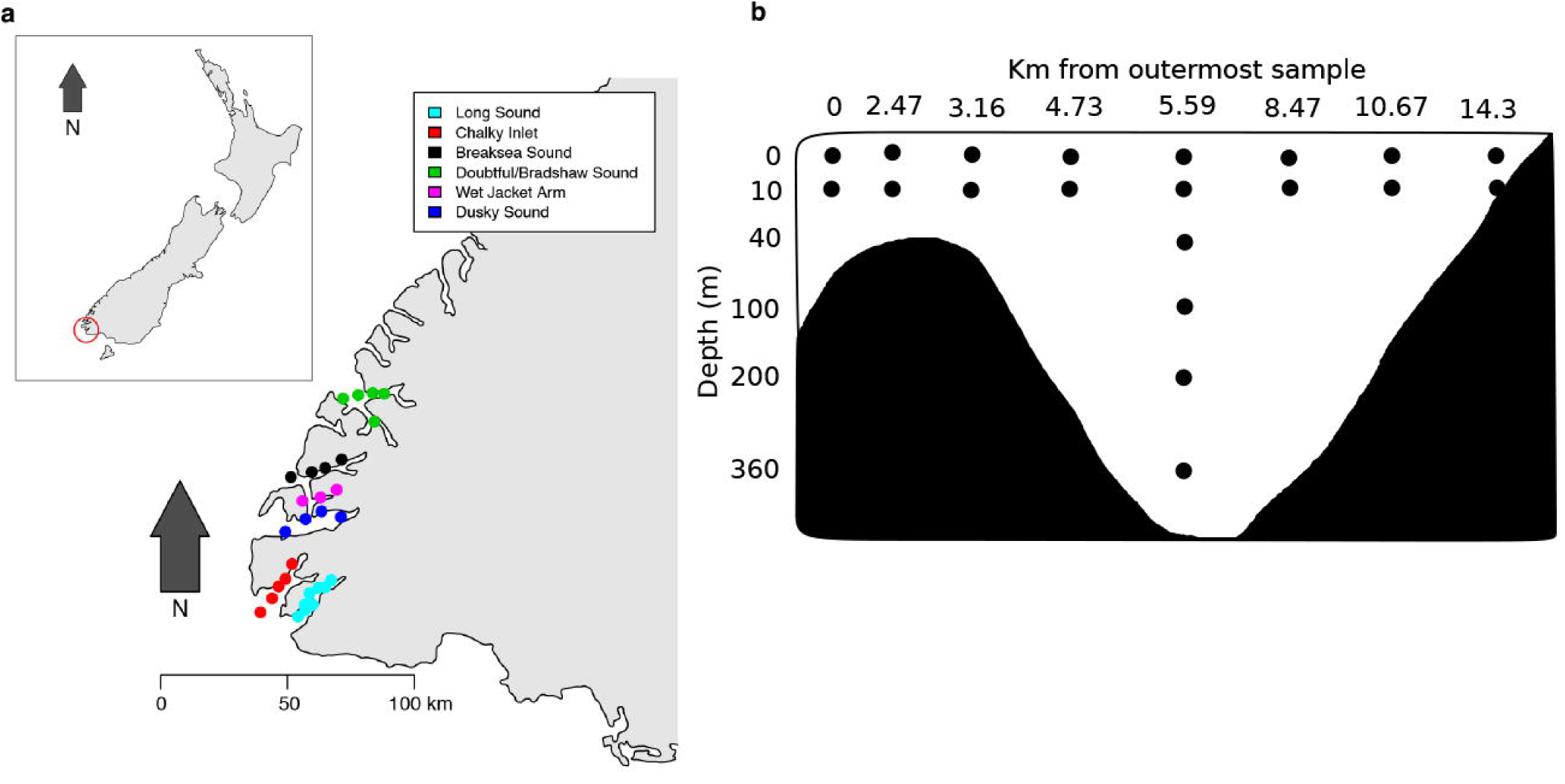
Changes in environmental parameters and significantly associated taxa along Long Sound’s horizontal axis. Environmental parameter changes were noted across Long Sound’s horizontal axis surface (a) and 10 m (b) in a discrete manner. Also shown is the mean relative abundance of phyla across Long Sound’s horizontal axis (c-h). Communities were based on 16S (left side) and 18S (right side) sequencing. The microbial community was correlated against distance from the outermost sample (c, d), salinity (e, f), temperature (g), and oxygen (h). Phyla <1% mean relative abundance have been identified and grouped into Rare Taxa (<1%). Error bars represent standard error of the mean abundance from repeated samplings. Mean relative abundance was calculated from pooled significantly correlated taxa.

When taxonomic changes where explored using the depth profile in Long Sound, large rearrangements were observed at the domain and all eukaryotic phyla level organisms in accordance to mixing regions (Fig. 6). At the surface, Eukaryota (mostly Opisthokonta) clearly dominated the community (>75%), but as depth increased the relative abundance of eukaryotes decreased within the first 40 m to <60%, reaching stable levels at depths > 100 m (<38%, mostly SAR). Concerning prokaryotes, Archaea were almost absent at the surface (<5%), increasing with depth, and stabilizing at around 40 m (>50%). In contrast, Bacteria remained largely unaffected by depth. At phylum level, changes in taxa abundance were only pronounced for some groups. The relative abundance of SAR increased with depth, while Opisthokonta decreased. The prokaryotic phyla Proteobacteria and Bacteroidetes, and the eukaryotic phyla SAR and Opisthokonta remained the most abundant taxa within Long Sound that significantly correlated with most tested environmental variables (Fig. 6, 7). Shifts in abundance for different groups where consistent within domains, but differed across them. Changes in prokaryotic taxa abundance occurred at 100-200 m, whereas eukaryotic shifts occurred much closer to the surface at 40 m.

**Figure 6.**
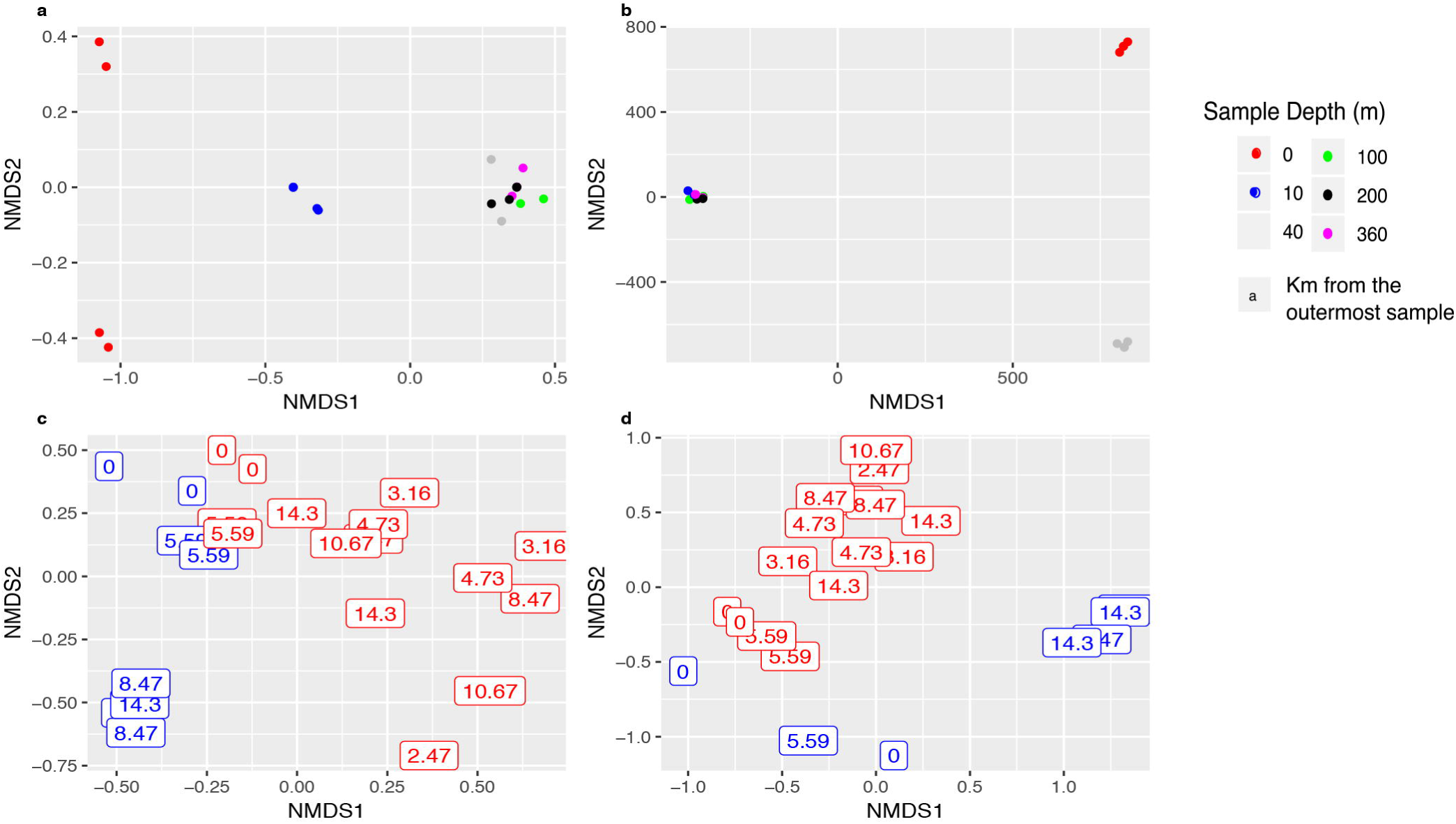
18S sequencing based microbial community of Long Sound’s vertical axis. Long Sound’s vertical community is shown at both the domain (a) and phyla (b) level as depth increases. Phyla <1% relative abundance have been identified and grouped into Rare Taxa (<1%). Mean relative abundance was calculated from pooled taxa.

**Figure 7.**
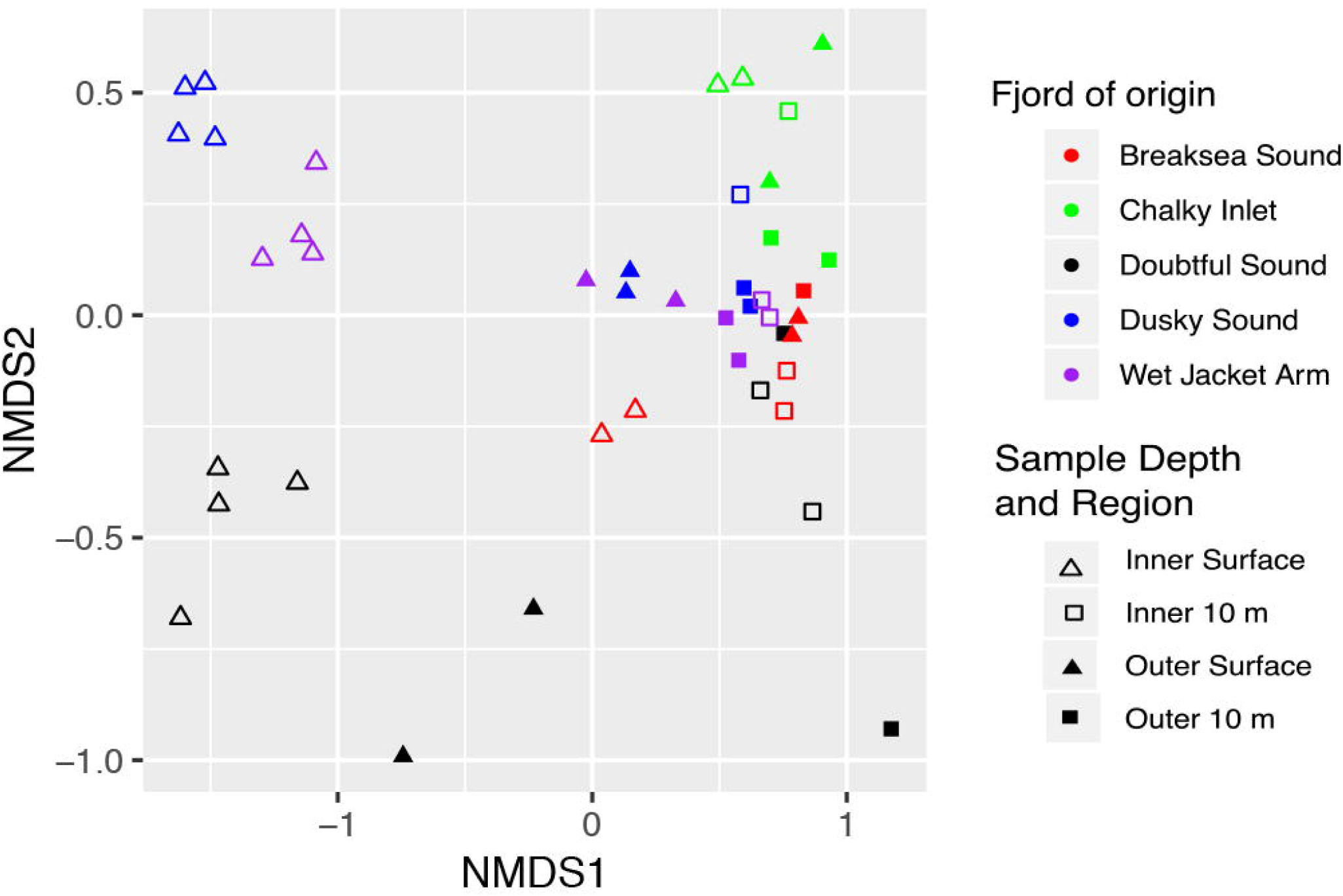
Changes in environmental parameters and significantly associated taxa along Long Sound’s vertical axis. Environmental parameter changes were noted across Long Sound’s vertical axis (a) in a discrete manner. Also shown is the mean relative abundance of phyla across Long Sound’s vertical axis. Communities were based on 16S (left side) and 18S (right side) sequencing. The microbial community was correlated against distance from the outermost sample (b, c), salinity (d, e), temperature (f), and oxygen (g. Phyla <1% mean relative abundance have been identified and grouped into Rare Taxa (<1%). Error bars represent standard error of the mean abundance from repeated samplings. Mean relative abundance was calculated from pooled significantly correlated taxa.

While some prokaryotes (i.e. Proteobacteria, Bacteroidetes, and Cyanobacteria), showed significant correlations with all tested environmental variables, this was not the case for eukaryotes (Fig. 5, 7). No eukaryotic taxa was found to significantly correlate with oxygen or temperature. However, we did see an inverse relationship for all Long Sound’s community patterns, most commonly between Proteobacteria and Bacteroidetes, and SAR and Opisthokonta (Fig. 5, 7).

## 4. Discussion

Our analyses indicate that a correlation exists between the diversity and community structure, and the depth and salinity conditions within six New Zealand fjords. These results are consistent with previous studies suggesting that changes in a fjord’s microbial diversity are primarily correlated with environmental conditions associated with depth, such as salinity (Bordalo and Vieira, 2005; Crump et al., 2004; Herlemann et al., 2011), light (Hölker et al., n.d.), organic matter (POC (Moncada et al., 2019), PON (Moncada et al., 2019), DOC (Li et al., 2012)), tannins (Baptist et al., 2008) and temperature (Berner et al., 2018; Hobbie, 1988). A LSL (low salinity layer), distinct from the lower marine layer (Citterio et al., 2017), is common across a fjord’s surface. Physicochemical differences between the marine and LSL layers could be responsible for large diversity and taxa shifts within the upper 10 m (Citterio et al., 2017). Hence, salinity could play a role for both the vertical and horizontal distribution of the community or act as a proxy for LSL mixing. Mixing zones, known to be particularly diverse due to hosting organisms from both environments (Gibbons and Gilbert, 2015), presented a particular strong response correlated with salinity changes (Cloern et al., 2017). The observed shift in salinity could most likely be associated with Fiordland’s characteristic extremely high rainfall levels and thus large freshwater inputs (NIWA, 2016) resulting in large differences in environmental conditions between the surface and the 10 m depth strata. Shifts in salinity, combined with high tannin concentrations, which limit light penetration to approximately 50 m (Nelson et al., 2002), could influence a rapid change in diversity patterns and taxa between depths, especially in the first 10 m. In addition, the retention times of different water masses in the fjords differ enormously. While the surface water stays several hours in the fjord and flows seawards constantly, the seawater below is slowly entrained staying several months in the fjord (Goebel et al., 2005) depending on freshwater inputs. The relatively shallow nature of the terminal sills at the fjord entrances (30 to 150 m (Stanton and Pickard, 1981)) limits the depth-associated effects by limiting the maximum salinity of the marine water within the fjord, since only the relatively shallow (and thus less saline) marine water enters and flows up the fjord (Pickrill, 1993; Stigebrandt, 2012; Talley et al., 2011), resulting in a less saline mixing layer between the marine water and LSL that would be associated with oceanic deep water. The relationship between salinity and depth (Maciejewska and Pempkowiak, 2014), is sustained on the scale of 100’s of meters within the ocean (Cummins, 1991) rather than the smaller scale relationship found within lakes (Garcia et al., 2013). But overall, Fiordland contains more unique characteristics that increase the effect of depth rather than those that decrease depth’s effect on its community.

### 4.1. Alpha diversity patterns were driven by depth and salinity

Mixing due to currents, wind or semi-continuous external freshwater inputs (e.g. waterfalls along the fjord walls) are likely responsible for the constant microbial (both prokaryotes and eukaryotes) richness observed along Long Sound’s surface as well as the varying retention times between the surface and 10 m layer. Microbial richness at 10 m depth decreased towards the fjord’s mouth, possibly caused by the lower levels of mixing and higher water retention times within Long Sound. Oxygen concentration decreases at 10 m towards the fjord’s head. Oxygen being consumed during the heterotrophic respiration of organic matter and a lack of renewal from atmospheric exchange due to the LSL, which persistently overcaps the underlying seawater.

The microbial diversity pattern found at the surface does not display a clear progression related to horizontal location, while at 10 m a seeming region-like richness and taxa change is noted for both prokaryotes and eukaryotes (Fig. 2, 3), most likely due to the varying retention time (Goebel et al., 2005) and dilution effects. The relative longer retention time at 10 m (compared to the LSL) and less wind mixing could allow for the establishment of more complex communities, a community increasing in complexity and diversity as it ages (Donlan, 2002). While the surface exhibits similar richness levels across the horizontal axis, its high flow rate corresponding to a high rate of horizontal dispersal (Campbell and Kirchman, 2013), and the homogeneity of phytoplankton communities within the LSL (and associated heterotrophic community) (Goebel et al., 2005) prevent the establishment of strong horizontal structure in microbial communities.

Observed richness along the vertical Long Sound axis correlated significantly with salinity but not depth on microbial diversity, corroborating the strong influence of the surface freshwater layer, constantly fed by external freshwater sources. Neither temperature nor oxygen showed a correlation with richness patterns, even though these variables are commonly considered to be very closely associated with depth. Much like marine snow often correlates with depth. Marine snow comes from the sinking of surface organic matter and organisms towards greater depths (Mestre et al., 2018), as such it is possible for microbes growing at the surface to be found at unexpected depths in an environment unsuitable for their growth. Much like marine snow, the dispersal effect, based upon the fact that due to their small size microbes disperse over large distances (Foissner, 2006), allows for microbes to be found in environments unsuitable for their optimal growth.

When taxonomic richness is compared among the six fjords, Doubtful Sound had a much higher richness level within the surface layer of the inner fjord. This 40 km long fjord is characterised by a larger LSL produced by the Manapouri Power Station freshwater output (Gibbs et al., 2000; Nelson et al., 2002). The deeper and more persistent LSL could have resulted in the observed increased microbial richness. An elevated flow rate and strong separation between the LSL and the underlying seawater layer likely prevented migration of microbial communities to the 10 m layer at the fjord’s head, also potentially decreasing vertical fluxes associated with marine snow. The inner surface region of five of the fjords studied presented higher microbial taxonomic richness levels, in contrast with the trend observed in Long Sound. As such, the patterns observed among the six fjords demonstrated that while microbial richness patterns within these fjords were correlated with depth, it was not in accordance to our initial hypothesis. Overall, our results indicate that both prokaryotic and eukaryotic richness patterns within all tested fjords present a clear correlation with depth.

### 4.2. Depth effects drove community changes between sites

Long Sound’s prokaryotic and eukaryotic outermost communities presented more similarities amongst themselves than the innermost communities, possibly due to a smaller LSL in those outer regions and the effects of increased mixing on horizontal dispersal. The innermost communities at different depths were just as dissimilar to each other as they were to the outermost communities, indicating a clear effect of strong vertical gradients in environmental conditions linked to the presence of a deep and distinct LSL. The LSL is at its deepest at the fjord’s head where the main freshwater runoff is located, and in Long Sound because of topographic isolation of the inner fjord basin the LSL is thick and has a higher retention time than in the other fjords. Towards the outermost regions, across the narrow passage that connects Long Sound with the outer coast Preservation Inlet, the LSL becomes much shallower, at times disappearing (Wing, 2009) and is subject to advection and mixing by ocean waves. As such, the outermost communities at both the surface and 10 m exist in non-LSL-influenced marine water. This contrasts with the communities closer to the fjord’s head, where the surface community is located primarily within the LSL, while the community at 10 m is found within LSL-influenced marine water. This degree of marine influence was also seen on the prokaryotic and eukaryotic vertical Long Sound axis NMDS plots. Grouping of communities into three separate layers was found, possibly due to depth associated environmental condition changes, specifically salinity and oxygen. However, prokaryotic communities of different depths were less dissimilar compared to eukaryotic communities. Both prokaryotes and eukaryotes showed that depth differences, and thus degree of marine influence, resulted in different communities. Based on this we hypothesised that each fjord would contain its own unique microbial community associated with the distinct environments within each fjord (e.g. depth, LSL depth, freshwater input, fjord length, entrance depth, etc.). Non-significant fjord clustering was noted for prokaryotes, the inner surface samples all clustering to the left of other samples from the same fjord, possibly due to the freshwater input carrying soil organisms from the fjord’s head. The separation of the inner surface samples reflects the increase in richness of the five-fjords but not the Long Sound richness patterns found in our study. Interestingly, the amount of freshwater appeared to influence the five-fjord NMDS, since Chalky Inlet (containing the smallest LSL) and Doubtful Sound (with the largest LSL) were located at opposite sides of the NMDS plot. Overall, diversity patterns matched with depth associated environmental changes across Long Sound. However, those same trends were not visible within the five-fjord diversity and community patterns.

### 4.3. Fiordland fjords community composition changes are mainly due to the effects depth has on environmental variables

Our study was the first to simultaneously investigate patterns in both prokaryotic and eukaryotic community diversity within fjords. To date, very few studies have focused on diversity patterns in fjordic environments, particularly those of the eukaryotic community, often excluding large taxa and targeting specific groups such as protists (Orsi et al., 2012; Piquet et al., 2010). The results obtained on dominantly identified phyla in the present study are consistent with previous reports on marine and fjord systems, which were dominated by Proteobacteria (Aldunate et al., 2018; Sinha et al., 2017; Spietz et al., 2015; Vander Roost et al., 2018; Zaikova et al., 2010), Bacteroidetes (Aldunate et al., 2018; Fernández-Gómez et al., 2013; Sinha et al., 2017; Spietz et al., 2015), Opisthokonta (Del Campo et al., 2015), and SAR (Guillou et al., 2008).

Proteobacteria include many different phylogenetic groups capable of diverse functional ability, from consuming dissolved organic matter (Wagner et al., 2006) to Sulphur oxidation (Frigaard et al., 2006; Yamamoto and Takai, 2011). Marine associated Bacteroidetes degrade particulate organic matter, containing significantly more proteases than non-marine Bacteroidetes (Fernández-Gómez et al., 2013). Marine associated Bacteroidetes attachment to phytoplankton may thus influence Bacteroidetes distribution, as has been previously observed within blooms (Pinhassi et al., 2004; Smith et al., 2017; Teeling et al., 2012). Cyanobacteria, a photoautotrophic oxygen producing prokaryote phyla (Hamilton et al., 2016), showed high abundance at the surface but was mostly absent at greater depths. The observed pattern is consistent with those found in other marine systems, abundance decreasing with light penetration and availability (Cantonati et al., 2014).

Opisthokonta are a diverse group of eukaryotes including organisms from both the animal and fungal kingdoms (Adl et al., 2018; Shalchian-Tabrizi et al., 2008), making it hard to determine the role they play within marine systems. Key characteristics of Opisthokonta include synthesis of extracellular chitin, cyst/spore/cell walls of filamentous growth, hyphae, and the extracellular digestion of substrates (Adl et al., 2018). Only 5% of SAR are from well sampled families making them a poorly understood phyla (Grattepanche et al., 2018). SAR is made up of Stramenopila, Alveolate, and Rhizaria. Stramenopila are mostly made up of autotrophic primary CO_2_ producers. A shared ecological function has not yet been determined for Alveolate, which made up the majority of the SAR distribution patterns (Table S2-S4). Alveolata are characterised by the presence of cortical alveoli (membrane-bound sacs underlying the cell membrane). Rhizaria is made up of a diverse range of free-living heterotroph lineages. Photosynthetic Rhizaria have also undergone photosynthetic endosymbiosis. But difficulty in isolation and cultivation makes Rhizaria and other SAR clades hard to study within laboratory settings.

Prokaryotes and eukaryotes shifted at different depths along vertical axis of Long Sound. While the prokaryotic community shifted at 100 m depth, the eukaryotic community shifted much closer to the surface at 40 m depth. The late shift of the prokaryotic community is most likely due to the presence of a permanent sulphur mat at ~150 m (James et al., 2018), while the eukaryotic community shift could be explained by the effects of light penetration, due to the high tannin concentration in the water due to the land runoff in periods of high rainfall. As such, even changes across the horizontal axis can be attributed to the differences in environmental factors derived from the effects of depth.

The environmental factors of depth, horizontal location, and salinity consistently correlated with both prokaryotic and eukaryotic diversity patterns. Oxygen and temperature did not match the taxa distribution pattern as closely as depth, horizontal location, and salinity. Thus, we hypothesise that oxygen and temperature were not as important as the other environmental variables in determining taxonomic diversity patterns. Indeed, depth and salinity, which always significantly correlated with community diversity, are likely much more influential in determining microbial community assemblages established within a fjord.

## 5. Conclusions

For the first time depth and salinity have been shown to act as major factors on both prokaryotic and eukaryotic diversity, community composition, and taxa patterns across and within fjords. However, our exploratory study did not focus on individual organisms or their interactions within the ecosystem. We instead focused on high taxa rank distributions due to environmental parameters; prompted by the lack of similar studies. Subsequent studies should focus on functional analyses across depths and individual interactions across the ecosystem. A high-resolution sampling of all fjords should be utilised instead of a region-based analysis. Regional analyses allow the identification of general trends along a fjord but lack the power to assess individual fjords. Unique fjord characteristics (e.g. additional freshwater sources) make direct cross-fjord studies difficult; cross-fjord studies should instead assess conserved trends across multiple fjords. Further refining our understanding of microbial communities within a fjord and the ecological reasoning behind their distribution.

## Acknowledgments

We thank the officers and crew of *RV Polaris II* and science staff involved. We also thank M. Meyers, B. Dagg and J. Wenley for their contribution to data collection. Federico Baltar was supported by a Rutherford Discovery Fellowship (Royal Society of New Zealand).

---

**Table S1.** Kruskal-Wallis tests on observed richness in relation to depth, distance, fjord of origin, salinity, oxygen, temperature for all studied fjords.

**Table S2.** Prokaryotic taxa significantly correlated with distance, depth, salinity, oxygen, and temperature along the horizontal axis of Long Sound fjord.

**Table S3.** Prokaryotic taxa significantly correlated with distance, depth, salinity, oxygen, and temperature along the vertical axis of Long Sound fjord.

**Table S4.** Eukaryotic taxa significantly correlated with distance, depth, salinity, oxygen, and temperature along the horizontal and vertical axis of Long Sound fjord.

**Table S5.** Mantel tests on fjord taxa in relation to depth, distance, fjord of origin, salinity, oxygen, temperature for all studied fjords.

## References

Adl, S.M., Bass, D., Lane, C.E., Lukeš, J., Schoch, C.L., Smirnov, A., Agatha, S., Berney, C., Brown, M.W., Burki, F., Cárdenas, P., Čepička, I., Chistyakova, L., del Campo, J., Dunthorn, M., Edvardsen, B., Eglit, Y., Guillou, L., Hampl, V., Heiss, A.A., Hoppenrath, M., James, T.Y., Karpov, S., Kim, E., Kolisko, M., Kudryavtsev, A., Lahr, D.J.G., Lara, E., Le Gall, L., Lynn, D.H., Mann, D.G., Massana i Molera, R., Mitchell, E.A.D., Morrow, C., Park, J.S., Pawlowski, J.W., Powell, M.J., Richter, D.J., Rueckert, S., Shadwick, L., Shimano, S., Spiegel, F.W., Torruella i Cortes, G., Youssef, N., Zlatogursky, V., Zhang, Q., Zhang, Q., 2018. Revisions to the Classification, Nomenclature, and Diversity of Eukaryotes. J. Eukaryot. Microbiol. 66, jeu.12691. https://doi.org/10.1111/jeu.12691

Aldunate, M., De la Iglesia, R., Bertagnolli, A.D., Ulloa, O., 2018. Oxygen modulates bacterial community composition in the coastal upwelling waters off central Chile. Deep. Res. Part II Top. Stud. Oceanogr. 1–12. https://doi.org/10.1016/j.dsr2.2018.02.001

Altschul, S.F., Madden, T.L., Schaffer, A.A., Zhang, J., Zhang, Z., Miller, W., Lipman, D.J., 1997. Gapped BLAST and PSI-BLAST: a new generation of protein database search programs. Nucleic Acids Res. 25, 3389–3402.

Arrigo, K.R., 2005. Marine microorgansisms and global nutrient cycles. Nature 437, 349–355. https://doi.org/10.1038/nature0415

Azam, F., Worden, A.Z., 2004. Microbes, Molecules, and Marine Ecosystems, in: Microbes, Molecules, and Marine Ecosystems. Science, pp. 1622–1625.

Baptist, F., Zinger, L., Clement, J.C., Gallet, C., Guillemin, R., Martins, J.M.F., Sage, L., Shahnavaz, B., Choler, P., Geremia, R., 2008. Tannin impacts on microbial diversity and the functioning of alpine soils: a multidisciplinary approach. https://doi.org/10.1111/j.1462-2920.2007.01504.x

Berdjeb, L., Parada, A., Needham, D.M., Fuhrman, J.A., 2018. Short-term dynamics and interactions of marine protist communities during the spring–summer transition. ISME J. 1–11. https://doi.org/10.1038/s41396-018-0097-x

Berner, C., Bertos-Fortis, M., Pinhassi, J., Legrand, C., 2018. Response of microbial communities to changing climate conditions during summer cyanobacterial blooms in the Baltic Sea. Front. Microbiol. 9. https://doi.org/10.3389/fmicb.2018.01562

Bokulich, N.A., Subramanian, S., Faith, J.J., Gevers, D., Gordon, J.I., Knight, R., Mills, D.A., Caporaso, J.G., 2013. Quality-filtering vastly improves diversity estimates from Illumina amplicon sequencing. Nat. Methods 10, 57–59. https://doi.org/10.1038/nmeth.2276

Bordalo, A.A., Vieira, M.E.C., 2005. Spatial variability of phytoplankton, bacteria and viruses in the mesotidal salt wedge Douro Estuary (Portugal). Estuar. Coast. Shelf Sci. 63, 143–154. https://doi.org/10.1016/j.ecss.2004.11.003

Brattegard, T., 1980. Why Biologists are interested in Fjords, in: Freeland, H.J., Farmer, D.M., Levings, C.D. (Eds.), Fjord Oceanography. Plenum Press, London, pp. 53–66. https://doi.org/10.1007/978-1-4613-3105-6

Campbell, B.J., Kirchman, D.L., 2013. Bacterial diversity, community structure and potential growth rates along an estuarine salinity gradient. ISME J. 7, 210–220. https://doi.org/10.1038/ismej.2012.93

Cantonati, M., Guella, G., Komárek, J., Spitale, D., 2014. Depth distribution of epilithic cyanobacteria and pigments in a mountain lake characterized by marked water-level fluctuations. Freshw. Sci. 33, 537–547. https://doi.org/10.1086/675930

Caporaso, J.G., Kuczynski, J., Stombaugh, J., Bittinger, K., Bushman, F.D., Costello, E.K., Fierer, N., Peña, A.G., Goodrich, J.K., Gordon, J.I., Huttley, G. a, Kelley, S.T., Knights, D., Koenig, J.E., Ley, R.E., Lozupone, C. a, Mcdonald, D., Muegge, B.D., Pirrung, M., Reeder, J., Sevinsky, J.R., Turnbaugh, P.J., Walters, W. a, Widmann, J., Yatsunenko, T., Zaneveld, J., Knight, R., 2010. correspondence QIIME allows analysis of highthroughput community sequencing data Intensity normalization improves color calling in SOLiD sequencing. Nat. Publ. Gr. 7, 335–336. https://doi.org/10.1038/nmeth0510-335

Caporaso, J.G., Lauber, C.L., Walters, W.A., Berg-Lyons, D., Huntley, J., Fierer, N., Owens, S.M., Betley, J., Fraser, L., Bauer, M., Gormley, N., Gilbert, J.A., Smith, G., Knight, R., 2012. Ultra-high-throughput microbial community analysis on the Illumina HiSeq and MiSeq platforms. ISME J. 6, 1621–1624. https://doi.org/10.1038/ismej.2012.8

Citterio, M., Sejr, M.K., Langen, P.L., Mottram, R.H., Abermann, J., Hillerup Larsen, S., Skov, K., Lund, M., 2017. Towards quantifying the glacial runoff signal in the freshwater input to Tyrolerfjord–Young Sound, NE Greenland. Ambio 46, 146–159. https://doi.org/10.1007/s13280-016-0876-4

Cloern, J.E., Jassby, A.D., Schraga, T.S., Nejad, E., Martin, C., 2017. Ecosystem variability along the estuarine salinity gradient: Examples from long-term study of San Francisco Bay. Limnol. Oceanogr. 62, S272–S291. https://doi.org/10.1002/lno.10537

Crump, B.C., Hopkinson, C.S., Sogin, M.L., Hobbie, J.E., 2004. Microbial Biogeography along an Estuarine Salinity Gradient: Combined Influences of Bacterial Growth and Residence Time Microbial Biogeography along an Estuarine Salinity Gradient: Combined Influences of Bacterial Growth and Residence Time. Appl. Environ. Microbiol. 70, 1494–1505. https://doi.org/10.1128/AEM.70.3.1494

Cummins, P.F., 1991. The deep water stratification of ocean general circulation models. Atmos. Ocean 29, 563–575. https://doi.org/10.1080/07055900.1991.9649417

Currey, R.J.C., Dawson, S.M., Slooten, E., Schneider, K., Lusseau, D., Boisseau, O.J., Haase, P., Williams, J.A., 2009. Survival rates for a declining population of bottlenose dolphins in Doubtful Sound, New Zealand: an information theoretic approach to assessing the role of human impacts. Ann. Zool. Fennici 19, 658–670. https://doi.org/10.1002/aqc

Del Campo, J., Mallo, D., Massana, R., de Vargas, C., Richards, T.A., Ruiz-Trillo, I., 2015. Diversity and distribution of unicellular opisthokonts along the European coast analysed using high-throughput sequencing. Environ. Microbiol. 17, 3195–207. https://doi.org/10.1111/1462-2920.12759

Donlan, R.M., 2002. Biofilms: microbial life on surfaces. Emerg. Infect. Dis. 8, 881–90. https://doi.org/10.3201/eid0809.020063

Edgar, R.C., 2010. Search and clustering orders of magnitude faster than BLAST. Bioinformatics 26, 2460–2461. https://doi.org/10.1093/bioinformatics/btq461

Falkowski, P.G., Fenchel, T., Delong, E.F., 2008. The Microbial Engines That Drive Earth ‘s Biogeochemical Cycles. Science (80-.). 320, 1034–1039. https://doi.org/10.1126/science.1153213

Fernández-Gómez, B., Richter, M., Schüler, M., Pinhassi, J., Acinas, S.G., González, J.M., Pedrós-Alió, C., 2013. Ecology of marine bacteroidetes: A comparative genomics approach. ISME J. 7, 1026–1037. https://doi.org/10.1038/ismej.2012.169

Foissner, W., 2006. Biogeography and dispersal of micro-organisms: A review emphasizing protists. Acta Protozool. 45, 111–136. https://doi.org/10.1.1.465.6672

Frigaard, N.U., Martinez, A., Mincer, T.J., DeLong, E.F., 2006. Proteorhodopsin lateral gene transfer between marine planktonic Bacteria and Archaea. Nature 439, 847–850. https://doi.org/10.1038/nature04435

Garcia, S.L., Salka, I., Grossart, H.P., Warnecke, F., 2013. Depth-discrete profiles of bacterial communities reveal pronounced spatio-temporal dynamics related to lake stratification. Environ. Microbiol. Rep. 5, 549–555. https://doi.org/10.1111/1758-2229.12044

Gibbons, S.M., Gilbert, J.A., 2015. Microbial diversity--exploration of natural ecosystems and microbiomes. Curr. Opin. Genet. Dev. 35, 66–72. https://doi.org/10.1016/j.gde.2015.10.003

Gibbs, M.T., Bowman, M.J., Dietrich, D.E., 2000. Maintenance of near-surface stratification in Doubtful Sound, a New Zealand fjord. Estuar. Coast. Shelf Sci. 51, 683–704. https://doi.org/10.1006/ecss.2000.0716

Goebel, N.L., Wing, S.R., Boyd, P.W., 2005. A mechanism for onset of diatom blooms in a fjord with persistent salinity stratification. Estuar. Coast. Shelf Sci. 64, 546–560. https://doi.org/10.1016/j.ecss.2005.03.015

Graham, E.B., Knelman, J.E., Schindlbacher, A., Siciliano, S., Breulmann, M., Yannarell, A., Beman, J.M., Abell, G., Philippot, L., Prosser, J., Foulquier, A., Yuste, J.C., Glanville, H.C., Jones, D.L., Angel, R., Salminen, J., Newton, R.J., Bu⁏rgmann, H., Ingram, L.J., Hamer, U., Siljanen, H.M.P., Peltoniemi, K., Potthast, K., Bañeras, L., Hartmann, M., Banerjee, S., Yu, R.-Q., Nogaro, G., Richter, A., Koranda, M., Castle, S.C., Goberna, M., Song, B., Chatterjee, A., Nunes, O.C., Lopes, A.R., Cao, Y., Kaisermann, A., Hallin, S., Strickland, M.S., Garcia-Pausas, J., Barba, J., Kang, H., Isobe, K., Papaspyrou, S., Pastorelli, R., Lagomarsino, A., Lindstro⁏m, E.S., Basiliko, N., Nemergut, D.R., 2016. Microbes as Engines of Ecosystem Function: When Does Community Structure Enhance Predictions of Ecosystem Processes? Front. Microbiol. 7, 1–10. https://doi.org/10.3389/fmicb.2016.00214

Grattepanche, J.-D., Walker, L.M., Ott, B.M., Paim Pinto, D.L., Delwiche, C.F., Lane, C.E., Katz, L.A., 2018. Microbial Diversity in the Eukaryotic SAR Clade: Illuminating the Darkness Between Morphology and Molecular Data. BioEssays 40, 1700198.https://doi.org/10.1002/bies.201700198

Guillou, L., Viprey, M., Chambouvet, A., Welsh, R.M., Kirkham, A.R., Massana, R., Scanlan, D.J., Worden, A.Z., 2008. Widespread occurrence and genetic diversity of marine parasitoids belonging to Syndiniales (Alveolata). Environ. Microbiol. 10, 3349–3365. https://doi.org/10.1111/j.1462-2920.2008.01731.x

Hamilton, T.L., Bryant, D.A., Macalady, J.L., 2016. The role of biology in planetary evolution: cyanobacterial primary production in low□oxygen Proterozoic oceans. Environ. Microbiol. 18, 325. https://doi.org/10.1111/1462-2920.13118

Henriques, I.S., Alves, A., Tacão, M., Almeida, A., Cunha, Â., Correia, A., 2006. Seasonal and spatial variability of free-living bacterial community composition along an estuarine gradient (Ria de Aveiro, Portugal). Estuar. Coast. Shelf Sci. 68, 139–148. https://doi.org/10.1016/j.ecss.2006.01.015

Herlemann, D.P.R., Labrenz, M., Jürgens, K., Bertilsson, S., Waniek, J.J., Andersson, A.F., 2011. Transitions in bacterial communities along the 2000 km salinity gradient of the Baltic Sea. ISME J. 5, 1571–1579. https://doi.org/10.1038/ismej.2011.41

Hobbie, J.E., 1988. A comparison of the ecology of planktonic bacteria in fresh and salt water. Limnol. Oceanogr. 33, 750–764. https://doi.org/10.4319/lo.1988.33.4part2.0750

Hölker, F., Wurzbacher, C., Weißenborn, C., Monaghan, M.T., Holzhauer, S.I.J., Premke, K., n.d. Microbial diversity and community respiration in freshwater sediments influenced by artificial light at night. https://doi.org/10.1098/rstb.2014.0130

Hope, R.M., 2013. Rmisc: Ryan Miscellaneous. CRAN.

Jakacki, J., Przyborska, A., Kosecki, S., Sundfjord, A., Albretsen, J., 2017. Modelling of the Svalbard fjord Hornsund. Oceanologia 59, 473–495. https://doi.org/10.1016/j.oceano.2017.04.004

James, M., Hartstein, N., Giles, H., 2018. Assessment of ecological effects of expanding salmon farming in Big Glory Bay, Stewart Island-Part 2 Assessment of effects.

Junge, W., Clausen, J., Penner-Hahn, J.E., Yocum, C.F., Dau, H., 2006. Photosynthetic Oxygen Production. Science 312, 1470–1472. https://doi.org/10.1126/science.312.5779.1470b

Lahti, L., Shetty, S., Blake, T., Salojarvi J, 2017. Tools for microbiome analysis in R. Bioconductor.

Lê, S., Josse, J., Husson, F., 2008. {FactoMineR}: A Package for Multivariate Analysis. J. Stat. Softw. 25, 1–18. https://doi.org/10.18637/jss.v025.i01

Li, D., Sharp, J.O., Saikaly, P.E., Ali, S., Alidina, M., Alarawi, M.S., Keller, S., Hoppe-Jones, C., Drewes, J.E., 2012. Dissolved organic carbon influences microbial community composition and diversity in managed aquifer recharge systems. Appl. Environ. Microbiol. 78, 6819–28. https://doi.org/10.1128/AEM.01223-12

Logue, J.B., Findlay, S.E.G., Comte, J., 2015. Editorial: Microbial responses to environmental changes. Front. Microbiol. 6, 1–4. https://doi.org/10.3389/fmicb.2015.01364

Maciejewska, A., Pempkowiak, J., 2014. DOC and POC in the water column of the southern Baltic. Part I. Evaluation of factors influencing sources, distribution and concentration dynamics of organic matter**This study was supported by the Baltic-C/BONUS Plus EUFP6 Project, statutory activities o. Oceanologia 56, 523–548. https://doi.org/10.5697/oc.55-3.523

McMurdie, P.J., Holmes, S., 2013. Phyloseq: An R Package for Reproducible Interactive Analysis and Graphics of Microbiome Census Data. PLoS One 8. https://doi.org/10.1371/journal.pone.0061217

Mestre, M., Ruiz-González, C., Logares, R., Duarte, C.M., Gasol, J.M., Sala, M.M., 2018. Sinking particles promote vertical connectivity in the ocean microbiome. Proc. Natl. Acad. Sci. 115, E6799–E6807. https://doi.org/10.1073/pnas.1802470115

Moncada, C., Hassenrück, C., Gärdes, A., Conaco, C., 2019. Microbial community composition of sediments influenced by intensive mariculture activity. FEMS Microbiol. Ecol. 95. https://doi.org/10.1093/femsec/fiz006

Nelson, W.A., Villouta, E., Neill, K.F., Williams, G.C., Adams, N.M., Slivsgaard, R., 2002. Macroalgae of Fiordland, New Zealand. Tuhinga. NIWA, 2016. Annual Climate Summary 2015 [WWW Document].

Oksanen, J., 2008. Vegan: an introduction to ordination. Management 1, 1–10. https://doi.org/intro-vegan.Rnw 1260 2010-08-17 12:11:04Z jarioksa processed with vegan 1.17-6 in R version 2.12.1 (2010-12-16) on January 10, 2011.

Orsi, W., Song, Y.C., Hallam, S., Edgcomb, V., 2012. Effect of oxygen minimum zone formation on communities of marine protists. ISME J. 6, 1586–1601. https://doi.org/10.1038/ismej.2012.7

Pickrill, R.A., 1993. Sediment yields in Fiordland. J. Hydrol. (New Zealand) 31, 39–55.

Pinhassi, J., Sala, M.M., Havskum, H., Peters, F., Guadayol, Ò., Malits, A., Marrasé, C., 2004. Changes in bacterioplankton composition under different phytoplankton regimens. Appl. Environ. Microbiol. 70, 6753–6766. https://doi.org/10.1128/AEM.70.11.6753-6766.2004

Piquet, A.M.T., Scheepens, J.F., Bolhuis, H., Wiencke, C., Buma, A.G.J., 2010. Variability of protistan and bacterial communities in two Arctic fjords (Spitsbergen). Polar Biol. 33, 1521–1536. https://doi.org/10.1007/s00300-010-0841-9

Quast, C., Pruesse, E., Yilmaz, P., Gerken, J., Schweer, T., Yarza, P., Peplies, J., Glöckner, F.O., 2013. The SILVA ribosomal RNA gene database project: Improved data processing and web-based tools. Nucleic Acids Res. 41, 590–596. https://doi.org/10.1093/nar/gks1219

R Core Team, 2018. R: A Language and Environment for Statistical Computing.

Robinson, M.D., McCarthy, D.J., Smyth, G.K., 2009. edgeR: A Bioconductor package for differential expression analysis of digital gene expression data. Bioinformatics 26, 139–140. https://doi.org/10.1093/bioinformatics/btp616

RStudio Team, 2016. RStudio: Integrated Development Environment for R.

Sakami, T., Watanabe, T., Kakehi, S., Taniuchi, Y., Kuwata, A., 2016. Spatial variation of bacterial community composition at the expiry of spring phytoplankton bloom in Sendai Bay, Japan. Gene 576, 610–617. https://doi.org/10.1016/j.gene.2015.10.011

Shalchian-Tabrizi, K., Minge, M.A., Espelund, M., Orr, R., Ruden, T., Jakobsen, K.S., Cavalier-Smith, T., 2008. Multigene Phylogeny of Choanozoa and the Origin of Animals. PLoS One 3, e2098. https://doi.org/10.1371/journal.pone.0002098

Sinha, R.K., Krishnan, K.P., Kerkar, S., Divya David, T., 2017. Spatio-Temporal Monitoring and Ecological Significance of Retrievable Pelagic Heterotrophic Bacteria in Kongsfjorden, an Arctic Fjord. Indian J. Microbiol. 57, 116–120. https://doi.org/10.1007/s12088-016-0621-5

Smith, M.W., Herfort, L., Fortunato, C.S., Crump, B.C., Simon, H.M., 2017. Microbial players and processes involved in phytoplankton bloom utilization in the water column of a fast-flowing, river-dominated estuary. Microbiologyopen 6. https://doi.org/10.1002/mbo3.467

Smith, R.W., Bianchi, T.S., Allison, M., Savage, C., Galy, V., 2015. High rates of organic carbon burial in fjord sediments globally. Nat. Geosci. 8, 450–453. https://doi.org/10.1038/ngeo2421

Spietz, R.L., Williams, C.M., Rocap, G., Horner-Devine, M.C., 2015. A dissolved oxygen threshold for shifts in bacterial community structure in a seasonally hypoxic estuary. PLoS One 10, 1–18. https://doi.org/10.1371/journal.pone.0135731

Stanton, B.R., Pickard, G.L., 1981. Physical Oceanography of the New Zealand Fiords.

Stigebrandt, A., 2012. Hydrodynamics and Circulation of Fjords, in: Bengtsson, L., Herschy, R.W., Fairbridge, R.W. (Eds.), Encyclopedia of Lakes and Reservoirs. Springer Netherlands, Dordrecht, pp. 327–344. https://doi.org/10.1007/978-1-4020-4410-6_247

Storesund, J.E., Sandaa, R.A., Thingstad, T.F., Asplin, L., Albretsen, J., Erga, S.R., 2017. Linking bacterial community structure to advection and environmental impact along a coast-fjord gradient of the Sognefjord, western Norway. Prog. Oceanogr. 159, 13–30. https://doi.org/10.1016/j.pocean.2017.09.002

Sullam, K.E., Matthews, B., Aebischer, T., Seehausen, O., Bürgmann, H., 2017. The effect of top-predator presence and phenotype on aquatic microbial communities. Ecol. Evol. 7, 1572–1582. https://doi.org/10.1002/ece3.2784

Talley, L.D., Pickard, G.L., Emery, W.J., Swift, J.H., 2011. Typical Distributions of Water Characteristics. Descr. Phys. Oceanogr. 1–4. https://doi.org/10.1016/B978-0-7506-4552-2.10016-2

Teeling, H., Fuchs, B.M., Becher, D., Klockow, C., Gardebrecht, A., Bennke, C.M., Kassabgy, M., Huang, S., Mann, A.J., Waldmann, J., Weber, M., Klindworth, A., Otto, A., Lange, J., Bernhardt, J., Reinsch, C., Hecker, M., Peplies, J., Bockelmann, F.D., Amann, R., 2012. Substrate-Controlled Succession of Marine Bacterioplankton Populations Induced by a Phytoplankton Bloom. Science (80-.). 336, 608–611. https://doi.org/10.1126/science.278.5338.631

Vander Roost, J., Daae, F.L., Steen, I.H., Thorseth, I.H., Dahle, H., 2018. Distribution Patterns of Iron-Oxidizing Zeta- and Beta-Proteobacteria From Different Environmental Settings at the Jan Mayen Vent Fields. Front. Microbiol. 9, 3008. https://doi.org/10.3389/fmicb.2018.03008

Wagner, M., Nielsen, P.H., Loy, A., Nielsen, J.L., Daims, H., 2006. Linking microbial community structure with function: Fluorescence in situ hybridization-microautoradiography and isotope arrays. Curr. Opin. Biotechnol. 17, 83–91. https://doi.org/10.1016/j.copbio.2005.12.006

Walsh, E.A., Kirkpatrick, J.B., Rutherford, S.D., Smith, D.C., Sogin, M., D’Hondt, S., 2016. Bacterial diversity and community composition from seasurface to subseafloor. ISME J. 10, 979–989. https://doi.org/10.1038/ismej.2015.175

Werner, J.J., Zhou, D., Caporaso, J.G., Knight, R., Angenent, L.T., 2012. Comparison of Illumina paired-end and single-direction sequencing for microbial 16S rRNA gene amplicon surveys. ISME J. 6, 1273–1276. https://doi.org/10.1038/ismej.2011.186

Wickham, H., 2018. dplyr: A Grammar of Data Manipulation. CRAN.

Wickham, H., 2016. ggplot2: Elegant Graphics for Data Analysis. Springer-Verlag New York.

Wickham, H., 2011. The Split-Apply-Combine Strategy for Data Analysis. J. Stat. Softw. 40, 1–29.

Wing, S.R., 2009. Decadal-scale dynamics of sea urchin population networks in Fiordland, New Zealand are driven by juxtaposition of larval transport against benthic productivity gradients. Mar. Ecol. Prog. Ser. 378, 125–134. https://doi.org/10.3354/meps07878

Yamamoto, M., Takai, K., 2011. Sulfur metabolisms in epsilon-and gamma-proteobacteria in deep-sea hydrothermal fields. Front. Microbiol. 2, 1–8. https://doi.org/10.3389/fmicb.2011.00192

Ying, M., Yonghui, Z., Nianzhi, J., Yang, S., Ning, H., 2009. Vertical distribution and phylogenetic composition of bacteria in the Eastern Tropical North Pacific Ocean. Microbiol. Res. 164, 624–633. https://doi.org/10.1016/j.micres.2008.01.001

Zahran, H.H., 1999. Rhizobium-legume symbiosis and nitrogen fixation under severe conditions and in an arid climate. Microbiol. Mol. Biol. Rev. 63, 968–89, table of contents.

Zaikova, E., Walsh, D.A., Stilwell, C.P., Mohn, W.W., Tortell, P.D., Hallam, S.J., 2010. Microbial community dynamics in a seasonally anoxic fjord: Saanich Inlet, British Columbia. Environ. Microbiol. 12, 172–191. https://doi.org/10.1111/j.1462-2920.2009.02058.x

